# Dual RNase and β-lactamase activity of a single enzyme encoded in most Archaea

**DOI:** 10.1101/667907

**Authors:** Seydina M. Diene, Lucile Pinault, Nicholas Armstrong, Said Azza, Vivek Keshri, Saber Khelaifia, Eric Chabrière, Gustavo Caetano-Anolles, Jean-Marc Rolain, Pierre Pontarotti, Didier Raoult

**Author notes:** **Corresponding author:** Prof. Didier Raoult, **Address:** MEPHI, IHU-Mediterranee Infection, 19-21 Bd Jean Moulin, 13005 Marseille, France. **Email**.

## Abstract

β-lactams targeting the bacterial cell wall are not active on archaea. Here, we figure out that annotation of genes as β-lactamase in Archeae on the basis of homologous genes, initially annotated β-lactamases, is a remnant of the identification of the original activities of this group of enzymes, which in fact, have multiple functions including nuclease, ribonuclease, β-lactamase, or glyoxalase; which may specialized over time. We expressed a class B β-lactamase enzyme from *Methanosarcina barkeri* that digest penicillin G. Moreover, while a weak glyoxalase activity was detected, a significant ribonuclease activity on bacterial and synthetic RNAs was demonstrated. The β-lactamase activity was inhibited by a β-lactamase inhibitor (sulbactam), but its RNAse activity was not. This gene appears to has been transferred to the *Flavobacteriaceae* group including *Elizabethkingia* genus in which the expressed gene shows a more specialized activity toward resistance to tienanmicin but no glyoxalase activity. The expressed class C-like β-lactamase gene, also from *Methanosarcina sp*., shows also hydrolysis activity and was more closely related to DD-peptidase enzymes than known bacterial class C β-lactamases. Our findings highlight the requalification needness of annotated enzymes as β-lactamases and the specification overtime of multipotent enzymes in different ways in Archaea and bacteria.

## Introduction

Antibiotics are part of the microorganism’s arsenal in their struggle to master microbial ecosystems^1^. Most antibiotics are non-ribosomal peptides assembled by megaenzymes, the non-ribosomal peptide synthetases (NRPS) that have structural motifs which appear to be among the oldest of the living world^2,3^. In the case of β-lactams, enzymes named β-lactamases have been so far identified thanks to their hydrolyzing activities on this antibiotic family. However, annotation of genomes of multiple living species have shown that homologous sequences to these β-lactamases were present in most living organisms, including those for which there are no known β-lactam targets as seen in bacteria^4^. This is the case in human where eighteen genes annotated as metallo-β-lactamases have been identified since 1999, some of which, known as for their nuclease and/or ribonuclease activities, exhibited indeed a β-lactamase activity^5^. In fact, metallo-β-lactamase (MBL) enzymes are characterized by conserved motif (i.e. HxHxDH) and residues shared by all MBL fold superfamily proteins including β-lactamases, glyoxalase IIs, nucleases, ribonucleases, flavoproteins, and others^6,7^. The same is true for Archaea, in which two groups of β-lactamases are present in the majority of Archaea^4^, which are not, by nature, susceptible to β-lactams, in whom an alternative role of these enzymes may be suspected, in particular that of nuclease, ribonuclease, or glyoxalase. Specifically to Archaea, there is evidence of transfer event of class B β-lactamase to a single bacterial groups, i.e. the *Flavobacteriacaea* and especially *Elizabethkingia* genus, which has an atypical antibiotic resistance profile, including resistance to thienamycin by an enzymatic mechanism (through GOB and Bla enzymes)^8,9^. It is highly probable and without presuming that genes annotated as β-lactamases in Archaea actually supports β-lactamase activity next to other activities used by Archaea. Thus, the purpose of this work is to express identified archaeal class B and C β-lactamases to measure its different enzymatic activities.

## Results

Blast analysis of known bacterial β-lactamase genes such as class A (TEM-24, SHV-12), class B (VIM-2, NDM-1), class C (CMY-12, AAC-1), and class D (OXA-23, OXA-58) show no or insignificant results (% identity ≤ 24) against the NCBI archaeal database. However, as described, ancestral sequences are capable of detecting remote homologous sequences from published biological databases^10^. Consequently, using constructed phylogenetic trees (cf. suppl. figures) of the four bacterial β-lactamase classes, an ancestral sequence for each class was inferred. From the four inferred ancestral sequences, homologous sequences in the archaeal database were identified for the class B and C β-lactamases (**Suppl. fig. S1 and S2**). No significant hits were obtained for the class A and D.

### Archaeal Class B metallo-β-lactamase

An archaeal β-lactamase appeared highly conserved in several classes of archaea including *Archaeoglobi, Methanomicrobia, Methanobacteria, Thermococci, Methanococci, Thermoplasmata* and *Thermoprotei* (**fig. 1; Suppl. Table S1**)^11^. To evaluate these archaeal enzymes activity, the protein from *Methanosarcina barkeri* (gi|851225341; 213 aa; 25.5 kDa)(**fig. 1; Suppl. Table S1**) was experimentally tested. Protein alignment of this latter with known bacterial metallo-β-lactamase proteins reveals conserved motifs/amino acids including Histidine118 (His118), Aspartic acid 120 (Asp120), His196, and His263, markers of this metallo-β-lactamase class B as previously described^12^(**fig. S3**). Three-dimensional (3D) structure comparison of this enzyme with known and well characterized proteins in the Phyre2 investigator database reveals 100% of confidence and 94% of coverage with the crystal structure of the New Delhi metallo-β-lactamase 1 (NDM-1; Phyre2 ID: c3rkjA)(**Suppl. Table S2**). To evaluate these archaeal enzymes activity, the MetbaB protein from *Methanosarcina barkeri* was experimentally tested. As expected, this enzyme exhibits a significant hydrolysis activity on nitrocefin (**fig. 2A, 2B**) (with determined kinetic parameters k_cat_=18.2×10^-3^ s^-1^, K_M_=820 μM and resulting k_cat_/K_M_=22.19 s^-1^.M^-1^) and on penicillin G, when measuring its complete degradation toward a single metabolite i.e. benzyl penilloic acid within three hours (**fig. 2C**). As shown on **Suppl. Figure S4**, the MetbaB activity was also evaluated in different pH and was optimal on nitrocefin at pH 7. Furthermore, to confirm the β-lactamase activity of this enzyme, the combination of nitrocefin with β-lactamase inhibitor sulbactam (at 1 μg/mL) was tested. As shown in **Figure 2A** (column 4), in the presence of sulbactam, no degradation of the nitrocefin β-lactam could be detected, suggesting a complete inhibition of the archaeal β-lactamase enzyme. This neutralizing activity was confirmed microbiologically on a *Pneumococcus* strain highly susceptible to penicillin (MIC =0.012 μg/ml) and highly resistant to sulbactam (MIC =32 μg/ml). Indeed, bacteria could grow in the presence of 0.1 μg/ml of penicillin incubated with the archaeal β-lactamase, but not when sulbactam was added, suggesting an inhibition of penicillin G enzymatic digestion (**fig. 2D**).

**Figure 1:**
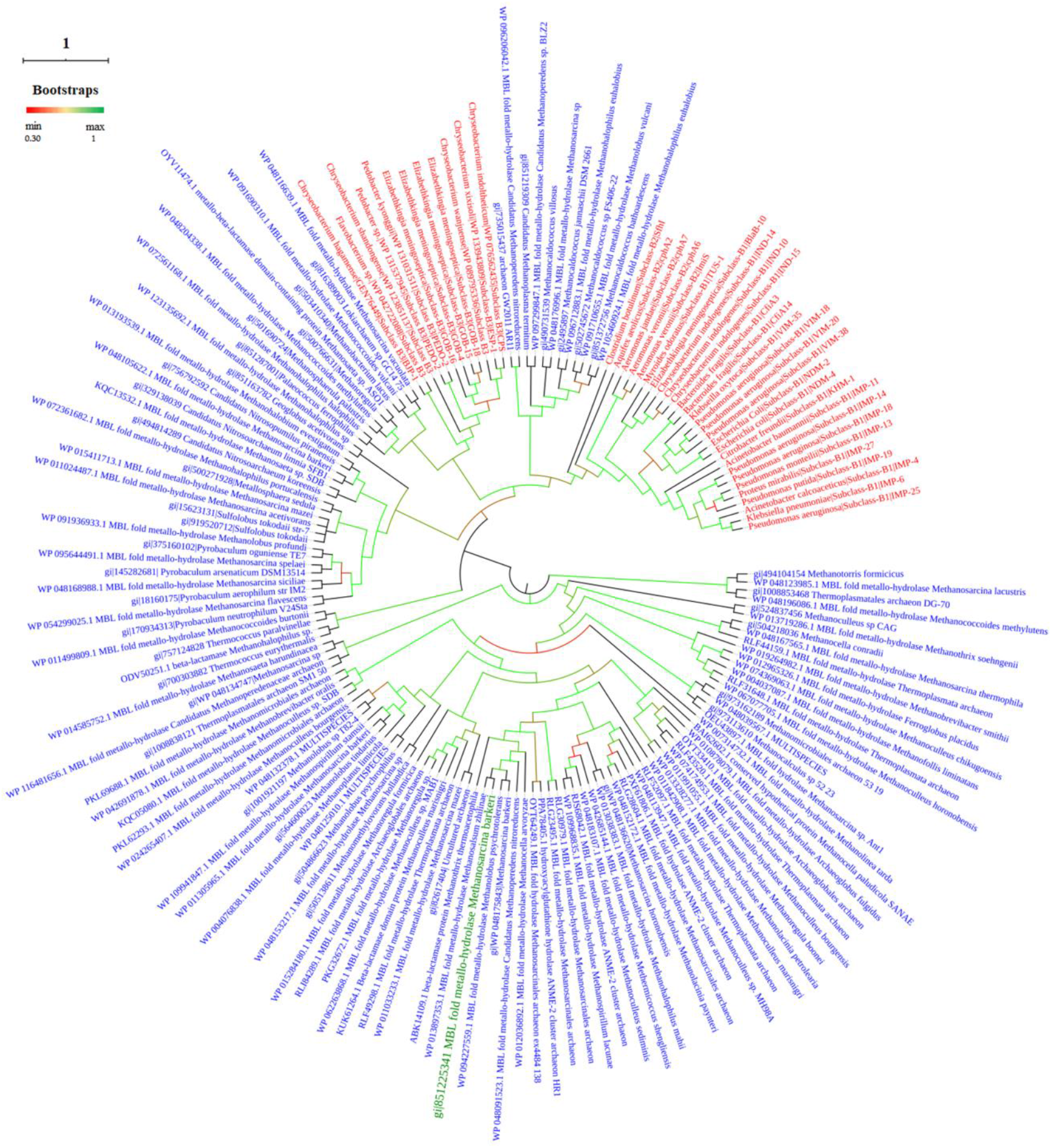
Phylogenetic Tree of Class B β-lactamases from archaea and bacteria. Archaeal sequence colored in green is which expressed and experimentaly tested. Bacterial β-lactamase sequences are colored in red whereas archaeal sequences are colored in bleue.

**Figure 2:**
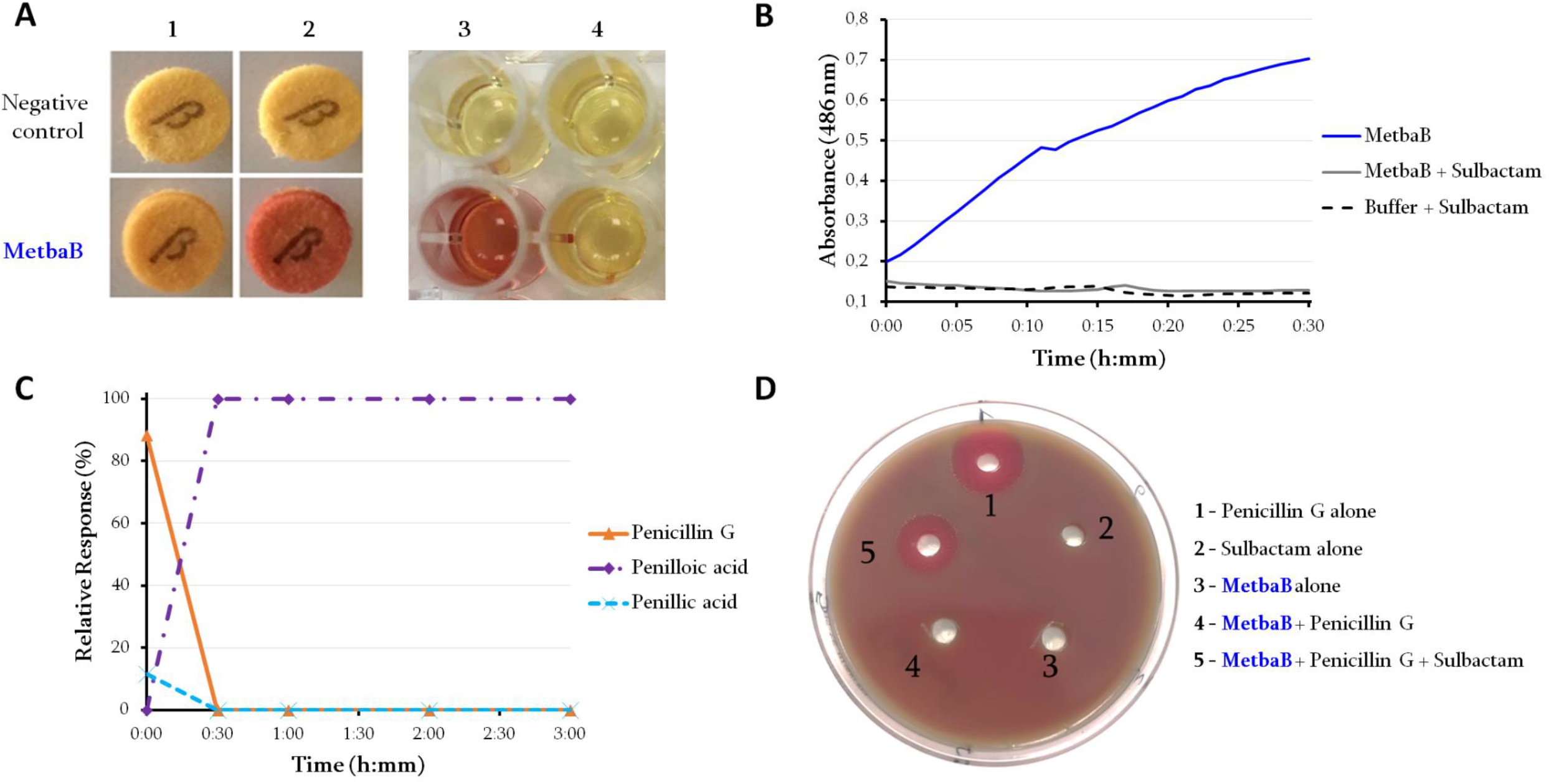
Characterization of the archaeal class B MBL (MetbaB) identified in *Methanosarcina barkerii*. **(A and B)**: β-lactamase activity of the *M. barkeri* Class B MBL enzyme (MetbaB) on a chromogenic cephalosporin substrate (Nitrocefin). **A1 and A2** refer to the nitrocefin degradation test using the BBL™ Cefinase™ paper disc respectively at t=0 and t=30 min. **A3** refers to this same test performed in liquid medium in the absence of sulbactam while **A4**, with the addition of 1 μg/ml sulbactam, both after 30 minutes of incubation; (**B**) monitored nitrocefin degradation by following the absorbance at 486 nm over time in the presence and absence of the β-lactamase inhibitor. **(C)**: LC/MS average relative response of screened metabolite compounds of penicillin G in the presence the *M. barkeri* Class B MBL enzyme monitored for three hours. Penicillin G (in orange) refers to the intact form of the antibiotic while penilloic acid (in purple) and penillic acid (in light blue) refer to the penicillin G metabolites. Penicillin G control in PBS did not show any degradation towards any metabolite (data not shown). **(D),** Microbiological test of the mixture of penicillin G (0.1 μg/ml) with the MetbaB enzyme in the presence and absence of sulbactam (15 μg/ml) on a *Pneumococcus* strain highly susceptible to penicillin G (MIC= 0.012 μg/ml) and highly resistant to sulbactam (MIC= 32 μg/ml). The halo around holes 1 and 5 reveals growth inhibition of the *Pneumococcus* strain. The absence of this halo around holes 2, 3, and 4 means no effect of the mixture on the *Pneumococcus* growth could be observed.

The antibiotic susceptibility testing of a recombinant *E. coli* mutant containing this Archaeal β-lactamase also revealed a reduced susceptibility to penicillin (from 1 μg/ml to 4 μg/ml) (data not shown). Interestingly, it appears that these Archaeal β-lactamases are closely related to bacterial enzymes known as “GOB” (AF090141), which are fully functional in vivo and present in a single bacterial family i.e. *Flavobacteriacaea*, especially in *Elizabethkingia* genus^8,13^ (**fig. 1 and suppl. fig. S5**). Indeed, we expressed the *bla*GOB-13 gene (AY647250) into *E. coli* BL21 strain and detected by LC-MS a full hydrolysis of imipenem by this enzyme through the accumulation of its metabolite (i.e. imipenemoic acid) over the time (**suppl. fig. S6**). As expected, the MetbaB enzyme hydrolyze also efficiency imipenem since its imipenemoic acid metabolite was detected after 24h (**suppl. fig S6**). Specific activities of GOB-13 and MetbaB enzymes were detected in the same order of magnitude on 1 mM of nitrocefin, which were 66 mU/mg and 24 mU/mg respectively.

However, the MBL protein sequences of this bacterial genus compared to those of Archaea reveal low similarities (less than 36%) and this therefore suggests an ancient HGT from an archaic phylum to this bacterial group, which furthermore exhibited natural β-lactam hydrolysis activity, previously considered to be fairly atypical for a bacterium (**Suppl. Table S3**). Thus, because archaea are resistant to ß-lactams, the role of these β-lactamases in these microorganisms may be the digestion of β-lactams to use it as a carbon source, as reported in the literature in bacteria^14^.

### Characterization of the DNAse and RNAse activities

As reported in the literature, MBL fold enzymes can have diverse functions such as nuclease, ribonuclease, and/or glyoxalase activities^6,7^ We tested here, the nuclease, ribonuclease, and glyoxalase activity of the expressed MetbaB enzyme. As presented on **Figure 3A**, while no nuclease activity on single and double-stranded DNAs was detected, extracted bacteria RNA (i.e. *E. coli* BL21 strain) was hydrolyzed by the archaeal MetbaB enzyme (**fig. 3B).** Moreover, using the RNaseAlert QC system kit able to detect unambiguously a real RNAse activity, we were able to confirm the MetbaB RNase activity with an average activity estimated to 0.359 mU/mg ± 0.107 (**fig. 3C**). Interestingly, in contrast, to β-lactamase activity, the MetbaB RNAse activity was not inhibited by the β-lactamase inhibitor, i.e. sulbactam (**Fig. 3B**).

**Figure 3:**
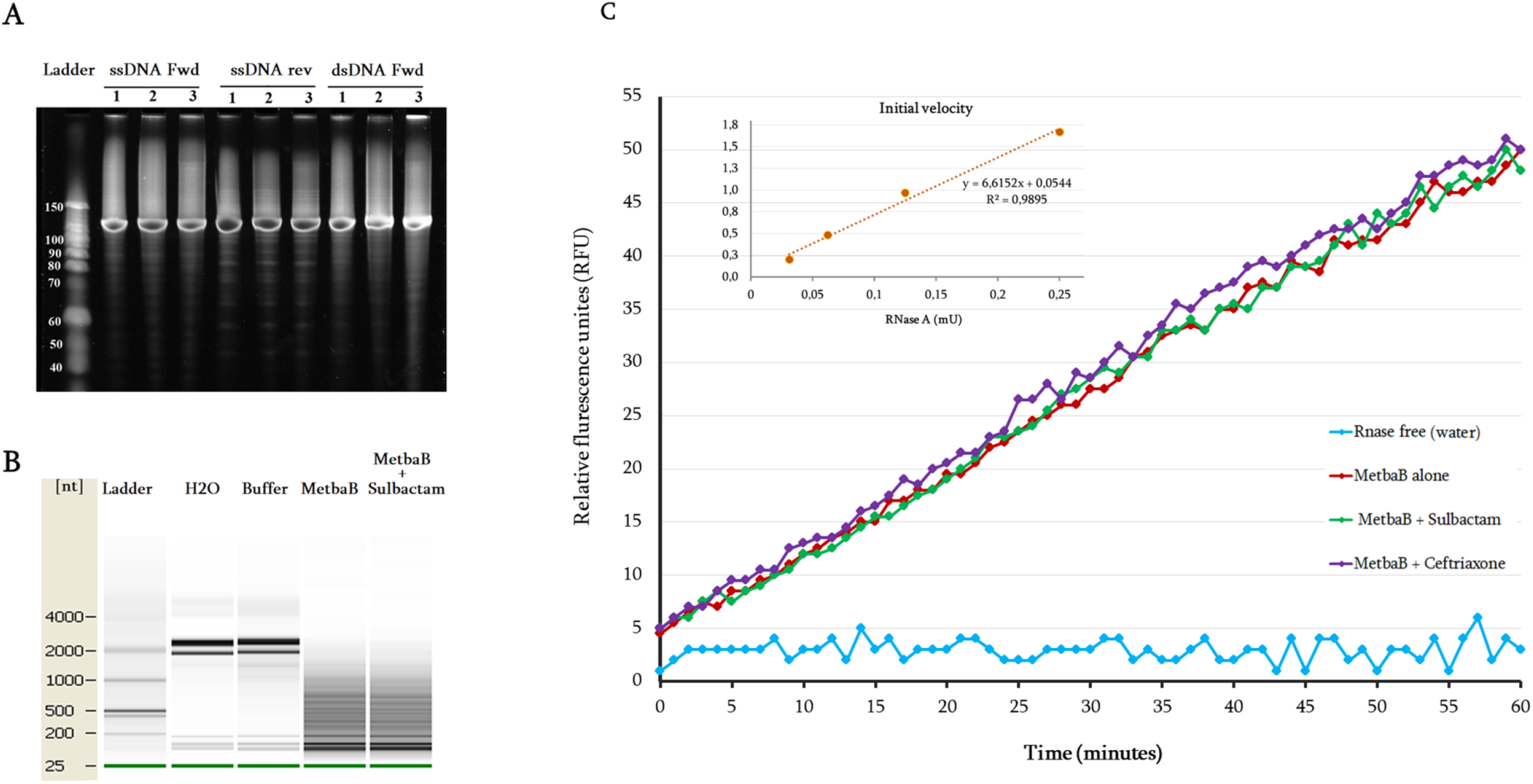
Evaluation of the DNAse and RNAse activity of the archaeal MetbaB enzyme. **(A)**: Effect of MetbaB enzyme on different synthetized DNA types (single and double-stranded DNA), each DNA type was tested here three time and no effect was observed. **(B)**: Effect of MetbaB enzyme on extracted bacterial RNA (*E. coli* BL21) in presence and absence of sulbactam (a β-lactamase inhibitor). The bacterial RNA was not degraded when incubated only with water and RNAse buffer. In contrast, this bacterial RNA was degraded when incubated with MetbaB enzyme alone or in presence of sulbactam. Presented gels in this figure are from different parts of the same gel **(C)**: confirmation of the RNAse activity of MetbaB enzyme using the RNAseAlert QC system kit. The RNAse activity is estimated according to the accumulated relative fluorescence during the time (here during one hour). The initial volacity of the enzyme was also determied.

### Glyoxalase activity

As presented on **Suppl. Figure S5**, the phylogenetic tree analysis shows that glyoxalase II sequences from bacteria and Eukarya appeared significantly related to archaeal MBL sequences. Base on that, the putative glyoxalase II activity of the MetbaB enzyme was then investigated. We were able to detect a weak activity of 3 mU/mg, using the Glyoxalase II activity kit from BioVision (Milpitas, CA, USA) (data not shown).

### Archaeal class C-like β-lactamases

Four significant sequences homologous to bacterial class C β-lactamase sequences were identified in archaea database using the inferred bacterial class C ancestor sequence (**fig. 4**; **Suppl. Table S1**). The phylogeny analysis shows that this third-class C-like of β-lactamases appears to be a very old class, a putative new clade, which cannot be identified without the reconstruction of the common ancestor (**fig. 4**). As shown in this figure, this class C-like enzyme appears more closely related to DD-peptidase enzymes than the known bacterial class C β-lactamases. Protein alignment reveals the same conserved and signature motifs (S^64^XXK and Y^150^XN) identified in bacterial class C β-lactamase (**Suppl. fig. S7**). Moreover, DD-peptidase enzymes can exhibit significant β-lactamase activity (10-fold higher β-lactams resistance) through punctual mutations in the coding sequence^15^. The three-dimensional (3D) structure comparison of this archaeal class C-like enzyme with known and well characterized proteins in the Phyre2 investigator database reveals 100% of confidence and 66% of coverage with the crystal structure of the octameric penicillin-binding protein (PBP) homologue from *pyrococcus abyssi* (Phyre2 ID: c2qmiH) (**Suppl. Table S2**). Similarly, the identified archaeal enzyme of this class C (gi|919167542) was also cloned in *E. coli* and found to be active in enzymatic level by hydrolyzing the nitrocefin (data not shown). This enzymatic activity was also confirmed by the kinetic assays showing the catalytic parameters kcat=9.67×10-3 s-1, Km=583.6 μM and kcat/ Km=16.57 s-1.M-1, according to Michaelis-Menten equation fitting (R^2^=0.984). However, the β-lactams susceptibility testing of the recombinant *E. coli* strains harboring this sequence reveals no reduced susceptibility as compared to the control *E. coli* strains.

**Figure 4:**
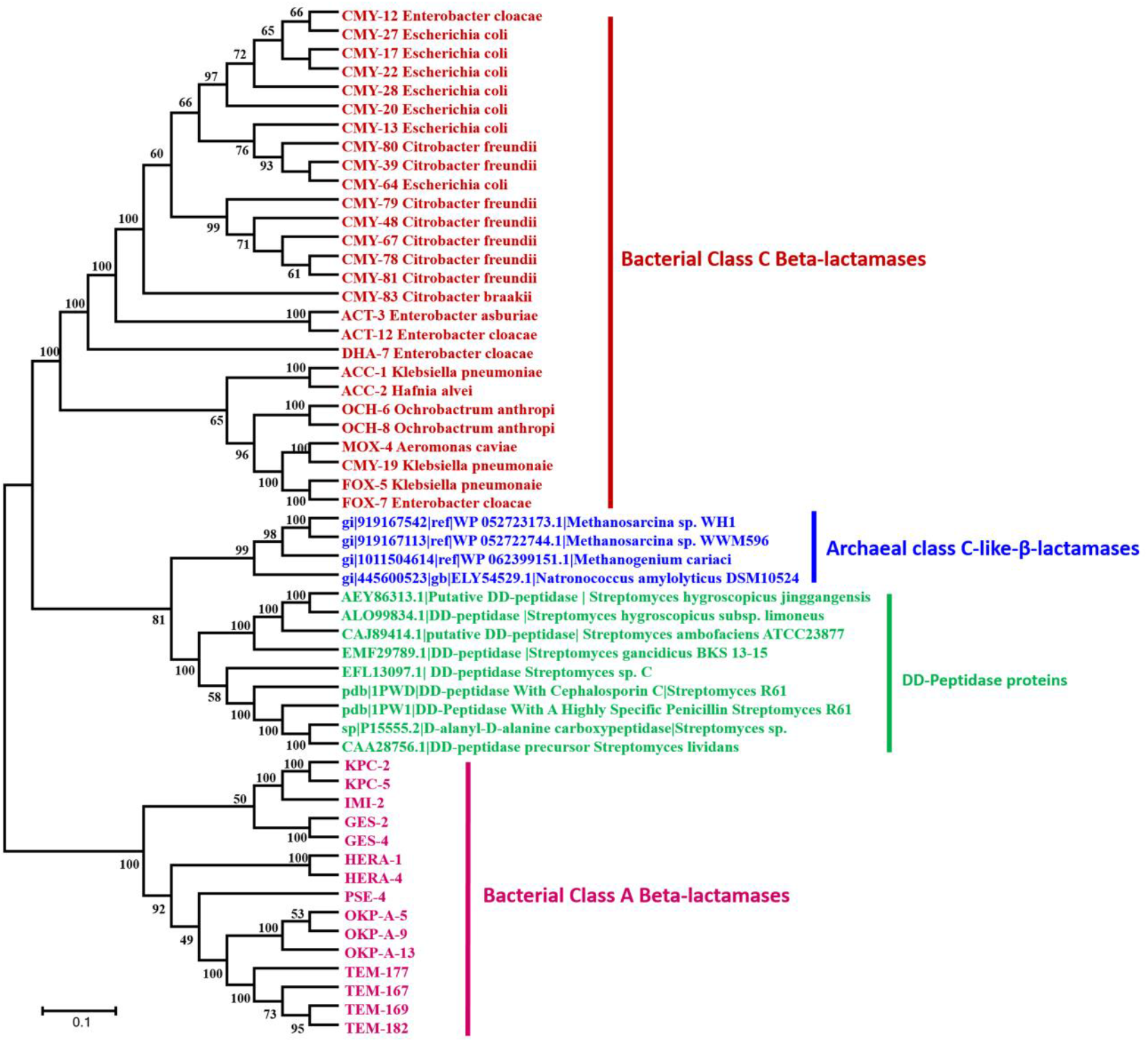
Phylogenetic Tree of Class C β-lactamases and DD-peptidases proteins (penicillin binding proteins). The class A β-lactamases is used as root.

## Discussion

In this study, we show that the activity of an Archaeal enzyme was triple including β-lactamase, ribonuclease, and glyoxalase and the annotation corresponding to only one of these activities is biologically unsatisfactory. The archaea microorganisms, in which the tested β-lactamase was identified (*Methanosarcina barkeri*), are fully resistant to all β-lactam antibiotics. This archaeal species has the largest genomes in the Archaea kingdom because of a massive horizontal gene transfer (HGT) from bacteria^16^. The identified class B β-lactamase sequences appear highly conserved and widespread in Archaea as previously reported using the hidden Markov model (HHM)-based profile^11^ and sequence transfer events have been observed into single bacterial family and particularly in *Elizabethkingia* genus, which has one of the largest spectrum of resistance to β-lactams known so far. In the current literature emerges evidence that β-lactamase enzymes, especially class B metallo-β-lactamase superfamily, have various activities such as β-lactamase, nuclease, ribonuclease, and glyoxalase^6,17,18^ (**Suppl. fig. S5**), which could justify the existence of enzymes in archaea acting also as β-lactamases. It has been reported that MBL enzyme superfamily exhibit a landscape of crossed activities, since each enzyme has on average 1.5 catalytic reactions in addition to its native activity^19^. Here, we were able to demonstrate that expressed archaeal β-lactamase enzyme (class B β-lactamase) can have triple activities (β-lactamase, ribonuclease, and glyoxalase). While the RNase activity was not inhibited by a β-lactamase inhibitor (sulbactam), the antibiotic hydrolyzing activity was inhibited by this inhibitor, a drug commonly used, in treatment of human infections, to inhibit bacterial β-lactamases^20^. The consequences of inhibiting the activities of these enzymes in the physiology of host organisms is an area that remain to be explored. On the basis of our findings, the role of β-lactamase like enzymes in Archaea appears not yet totally understood. There is confusion between the annotation of nucleases/ribonucleases and β-lactamase enzymes. Both activities can be conserved in archaea and this is likely to demonstrate the ancient origin of MBL nucleases first and secondly, the risk of false annotation from the first identified enzymatic activity of the newly identified enzymes. Our findings suggest that archaeal β-lactamases are as ancestral as those of bacteria, and HGT events have occurred from archaea to bacteria, where enzymes can be specialized to other roles for more efficiency, like observed in *Elizabethkingia*. As presented in **Suppl. Figure S8**, we purpose a putative evolution scenario of enzymatic activities of MBLs in which ancestor MBL sequence has best hit protein in PDB (Protein Database Bank), a glyoxalase II (1XM8_A) exhibiting two different metal ions (Fe and Zn) in his catalytic site. Its evolution over the time resulted to (i) archaeal MBLs, e.g. MetbaB which matches in PDB with a MBL superfamily fold (AZZI_A) exhibiting only Fe ions in his catalytic site and has different enzymatic activities as shown in our present study and (ii) bacterial GOB enzymes (K0W_A) bearing only Zn ions for an activity more specific and more efficient against β-lactam antibiotics.

Finally, the existence of enzymes in the world of archaea with multiple activities such as β-lactamase, ribonuclease, and glyoxalase, is showing that β-lactamase enzymes are not only a defense system against β-lactam antibiotics. The use of antibiotics as a nutriment sources for archaea as key to degrade β-lactam molecules and use them as carbon sources as described in bacteria, is a plausible hypothesis^14,21–23^.

## Materials and Methods

### Sequence analysis

A total of 1,155 amino acid sequences were retrieved (Class A: 620; B: 174; C: 151, and D: 210) from the ARG-ANNOT database^24^. The phylogenetic trees were inferred using the approximate maximum-likelihood method in FastTree^25^. For a detailed and comprehensive diversity analysis, a few sequences from each clade of the trees were selected as representatives of the corresponding clades (labeled in red in **Suppl. fig. S1 and S2**).

The ancestral sequence was inferred using the maximum-likelihood method conducted by MEGA6^26^ software. Then, these ancestral sequences were used as queries in a BlastP^27^ search (≥ 30% sequence identity and ≥ 50% query coverage) against the NCBI-nr archaeal database. For Class C β-lactamase analysis, DD-peptidase sequences (penicillin binding proteins) were downloaded from the NCBI database. 2515 sequences were selected for local Blast analysis with the archaeal Class C-like β-lactamase used as query sequence (gi| 919167542). From this analysis, 24 DD-peptidase sequences were identified as homologous to the query and thus used for further phylogenetic tree analysis. The selected archaeal sequences were aligned with known bacterial β-lactamase sequences (representative sequences of a known clade from the guide tree) using the multiple sequence alignment algorithm MUSCLE^28^ and the phylogenetic tree was inferred using FastTree^25^.

### Antibiotic susceptibility testing

The antibiotic susceptibility testing of two *Methanosarcina (barkeri* and sp.) isolates was performed on 15 antibiotics including ampicillin, ampicillin/sulbactam, penicillin, piperacillin, piperacillin/tazobactam, cefoxitin, ceftriaxone, ceftazidime imipenem, meropenem, aztreonam, gentamicin, ciprofloxacin, amikacin, and trimethoprim-sulfamethoxazole (I2a, SirScan Discs, France). A filtred aqueous solution of each antibiotic was prepared anaerobically in a sterilized Hungate tubes at concentration of 5 mg/ml. Then, 0.1 ml of each one of these solutions was added to a freshly inoculated culture tube containing 4.9 ml of the tested stain to obtain a final concentration of 100μg/ml for each antibiotic herein tested. The mixture of antibiotic and archaeal culture was then incubated at 37°C and the growth of archaea was observed after 5 to 10 days incubation depending on the tested strain. Control cultures without antibiotic were also incubated in the same conditions to assess the strain growth and non-inoculated culture tubes were used as negative control.

### In vitro assessment of the β-lactamase activity

#### Protein expression and purification

Genes encoding the selected β-lactamases including the *Methanosarcina* class B β-lactamase protein (gi|851225341), class C-like β-lactamase protein (gi|919167542) and the *Elizabethkingia* GOB-13 (AY647250) were synthesized by GenScript (Piscataway, NJ, USA) and optimized for protein expression in *Escherichia coli* in the pET24a(+) expression vector. The details of this protein expression and purification of the recombinant proteins are previously described^29^. Purified proteins were then subjected to β-lactamase activity detection as previously described using nitrocefin and penicillin in presence and absence of sulbactam^29^. Furthermore, the activity of MetbaB enzyme was evaluated at different pH (between pH7 and pH10) using the same nitrocefin assay conditions. The kinetic parameters of the MetbaB enzyme were determined using the same conditions as previously reported^29^. In addition to performed analyses described below, the β-lactam hydrolysis activity was monitored by Liquid Chromatography-Mass Spectrometry (LC-MS) on penicillin G and imipenem in presence and absence of the β-lactamase inhibitor, i.e. sulbactam as we described previously^29^.

#### Imipenem antibiotic degradation monitored by Liquid Chromatography-Mass Spectrometry (LC-MS)

A stock solutions at 10 mg/ml of Imipenem and Cilastatine was freshly prepared in water from the perfusion mixture of both compounds (500 mg/500 mg; Panpharma, Luitre, France). 100 μL of GOB-13 and MetbaB enzyme solutions at 1 mg/mL were spiked with Imipenem/Cilastatine at final concentrations of 10 μg/ml, then incubated at room temperature. Negative controls consisted of PBS spiked with Imipenem/Cilastatine. Triplicate samples were prepared and for each replicate 30 μL of solution was collected at 0, 4 and 24 hours. Then, 70 μL of acetonitrile was added to each sample, and tubes were vortexed 10 minutes at 16000 g to precipitate proteins. The clear supernatant was collected for analysis using an Acquity I-Class UPLC chromatography system connected to a Vion IMS Qtof ion mobility-quadrupole-time of flight mass spectrometer as previously described (**Reference**) with modifications. For each sample stored at 4°C, 10 μL was injected into a reverse phase column (Acquity BEH C18 1.7 μm 2.1×50 mm, Waters) maintained at 50°C. Compounds were eluted at 0.5 ml/min using water and acetonitrile solvents each containing 0.1% formic acid. The following composition gradient was used: 5% during 1 minute to70% acetonitrile, 95 % acetonitrile for a 1-minute wash step, and back to the initial composition for 1-minute. Compounds were ionized in the positive mode using a Zspray electrospray ion source with the following parameters: capillary/cone voltages 2.5/40V, and source/desolvation temperatures 120/450°C. Ions were then monitored using a High Definition MS(E) data independent acquisition method with the following settings: travelling wave ion mobility survey, 50-1000 m/z, 0.1 s scan time, 6 eV low energy ion transfer, and 20-40 eV high energy for collision-induced dissociation of all ions (low/high energy alternate scans). Mass calibration was adjusted within each run using a lockmass correction (Leucin Enkephalin 556.2766 m/z). The Vion instrument ion mobility cell and time-of-flight tube were calibrated beforehand using a Major Mix solution (Waters) to calculate collision cross section (CCS) values from ion mobility drift times and mass-to-charge ratios. 4D peaks, corresponding to a chromatographic retention time, ion mobility drift time and parents/fragments masses, were then collected from raw data using UNIFI software (version 1.9.3, Waters). As reported, the lactam ring of Imipenem can be hydrolyzed to form the Imipenemoic acid structure. A list of known chemical structures including Imipenem, Imipenemoic acid and Cilastatine were targeted with the following parameters: 0.1 minutes retention time window, 5 % CCS tolerance, 5 ppm m/z tolerance on parent adducts (H+ and Na+) and 10 mDa m/z tolerance on predicted fragments. Retention times and CCS values were previously measured from antibiotics standards in order to perform subsequent accurate structures screening (Imipenem: 0.3 minutes/169 Å^2^; Imipenemoic acid: 1.8 minutes/242 Å^2^). The MS Responses of Imipenem and Imipenemoic acid were normalized using the MS Response of Cilastatine (ratio) for data interpretation. Phase I chemical transformations were also screened against the raw data and showed that the hydrolysis (Imipenemoic acid) was an abundant metabolite in the case of GOB-13and MetbaB.

### DNAse and RNAse activity evaluation

To evaluate the DNAse activity of the MetbaB enzyme, synthesized single-stranded forward and reverse DNAs and double-stranded DNA of 130-bp were used as substrates. Double-stranded DNA was obtained by annealing forward and reverse single-stranded DNAs in thermocycler at temperatures decreasing from 95°C to 25°C during 1 h. Moreover, the RNAse activity was assessed using the RNaseAlert QC System kit (Fisher Scientific, Illkirch, France). This assay uses a fluorescence-quenched oligonucleotide probe as substrate that emits a fluorescent signal in the presence of RNase activity. Fluorescence was monitored continuously at 37°C for 1h by a Synergy HT plate reader (BioTek Instruments SAS, Colmar, France) with a 485/528 nm filter set. The RNase activity was then determined using supplied RNase A used as a standard (10 mU/mL). In addition to the tested RNaseAlert QC system kit, based on a synthetic RNA, purified *Escherichia coli* RNA using RNeasy columns (Invitrogen, Carlsbad, CA, USA) was also tested. Enzymatic reactions were performed by incubating each RNA samples with 15 μg of expressed MetbaB protein in Tris-HCl buffer 50 mM, pH 8.0, sodium chloride 0.3 M, using a final volume of 20 μL at 30°C for 2 h. After incubation, the material was loaded onto denaturing PolyAcrylamide Gel Electrophoresis (dPAGE) at 12% or analyzed using the Agilent RNA 6000 Pico LabChip kit on an Agilent 2100 Bioanalyzer (Agilent Technology, Palo Alto, CA, USA). Of course, RNase activities were assayed in the absence or presence of 10μg/mL of sulbactam. Negative controls were made with all used reagents (RNase free water, enzyme buffer) but also with bacterial culture without expression vector containing *metbaB* gene. Each experiment was performed at least in triplicate.

### Glyoxalase II Activity assay

Glyoxalase II (GloII) activity assays were performed using the Glyoxalase II Activity kit from BioVision (Milpitas, CA, USA) and monitored with a Synergy HT microplate reader (BioTek, Winooski, VT, USA). Reactions were carried out in triplicate at room temperature in a 96-well plate with a final volume of 100 μL for each well. Degradation of the GloII substrate was monitored for 40 minutes following absorbance variations at 450 nm, corresponding to the production of D-Lactate that reacts with a chromophore provided in the reaction mix. A D-Lactate Standard curve was plotted and allowed quantification of produced D-Lactate with our enzyme and calculation of its specific activity.

## Acknowledgements

Financial support from the IHU Mediterranee Infection, Marseille France and American Journal Experts (AJE) for English corrections of the manuscript are gratefully acknowledged.

## Author contributions

Didier Raoult. conceived and designed the study. Seydina M. Diene, Lucile Pinault, Vivek Keshri, Nicholas Armstrong, Said Azza, Saber Khelaifia, Eric Chabrière, Jean-Marc Rolain, Pierre Pontarotti, and Didier Raoult analysed and interpreted data. Seydina M. Diene, Lucile Pinault, Vivek Keshri, Nicholas Armstrong, Said Azza, Gustavo Caetano-Anolles, Eric Chabrière, Jean-Marc Rolain, Pierre Pontarotti, and Didier Raoult drafted the manuscript and/or made critical revisions. All of the authors read and approved the final manuscript.

