## Supplementary Figures and Tables for "Dual RNase and β-lactamase activity of a single enzyme encoded in most Archaea"

1. Aix Marseille Univ., MEPHI, IHU-Méditerranée Infection, Marseille, France.
2. Assistance Publique-Hôpitaux de Marseille (AP-HM), IHU-Méditerranée Infection, Marseille, France.
3. IHU-Méditerranée Infection, Marseille, France.
4. Evolutionary Bioinformatics Laboratory, Department of Crop Sciences, University of Illinois at Urbana-Champaign, Urbana, IL 61801, USA.
5. CNRS, Marseille, France.

**Suppl. Table S1:** Best blast Hits of ancestor β-lactamase sequence against the NCBI Archaea database

| Query id | NCBI hit gi | Organism hits | Size (bp) | Descriptions | % Identity | E-value | % Q_Cov |
|---|---|---|---|---|---|---|---|
| | gi\|564600023 | *Methanolobus tindarius* | 639 | Zn-dependent hydrolase | 34.62 | 5.26E-09 | 51.17 |
| | gi\|503410340 | *Methanobacterium lacus* | 645 | Hydrolase | 34.23 | 0.018 | 57.75 |
| | gi\|504218036 | *Methanocella conradii* | 624 | Hydrolase glyoxylase | 33.58 | 1.17E-06 | 53.99 |
| | gi\|145282681 | *Pyrobaculum arsenaticum* DSM 13514 | 621 | Beta-lactamase domain protein | 33.33 | 2.14E-07 | 52.58 |
| | gi\|375160102 | *Pyrobaculum oguniense* TE7 | 621 | Zn-dependent hydrolase glyoxylase | 32.52 | 8.80E-05 | 52.58 |
| | gi\|2495897 | *Methanocaldococcus jannaschii* DSM2661 | 618 | Probable metallo-hydrolase MJ0296 | 32.17 | 1.76E-04 | 50.70 |
| | gi\|1008838121 | *Thermoplasmatales archaeon* SM1-50 | 651 | Hypothetical protein | 32.14 | 0.28 | 54.46 |
| **Ancestor class B β-lactamase** | gi\|82617404 | uncultured archaeon | 645 | Hypothetical protein | 32.12 | 1.82E-05 | 53.05 |
| | gi\|1008853468 | *Thermoplasmatales archaeon* DG-70 | 606 | Hypothetical protein | 31.78 | 0.011 | 51.17 |
| | gi\|851372756 | *Methanocaldococcus bathoardescens* | 567 | MBL fold metallo-hydrolase | 31.58 | 1.05E-04 | 50.70 |
| | **gi\|851225341** | ***Methanosarcina barkeri*** | **641** | **MBL fold hydrolase (MetbaB)** | **31.54** | **4.88E-05** | **51.17** |
| | gi\|851312700 | *Methanosarcina barkeri* | 641 | MBL fold hydrolase | 31.54 | 2.85E-05 | 51.17 |
| | gi\|490731539 | *Methanocaldococcus villosus* | 540 | Hypothetical protein | 31.53 | 1.87E-05 | 50.70 |
| | gi\|1001921197 | *Methanolobus* sp. T82-4 | 672 | Hypothetical protein AWU59_1880 | 31.39 | 2.10E-06 | 53.99 |
| | gi\|505138611 | *Methanomethylovorans hollandica* | 645 | Zn-dependent hydrolase glyoxylase | 31.39 | 6.56E-08 | 54.46 |
| | gi\|501690724 | *Methanosphaerula palustris* | 696 | MBL fold hydrolase | 31.37 | 0.002 | 56.34 |

| | | | | | | |
|---|---|---|---|---|---|---|
| gi\|18160175 | *Pyrobaculum aerophilum* str. IM2 | 624 | Possibly metallo-beta-lactamase superfamily | 30.89 | 2.21E-06 | 52.58 |
| gi\|494814289 | *Candidatus Nitrosoarchaeum koreensis* | 611 | Zn-dependent hydrolase | 30.83 | 0.029 | 54.46 |
| gi\|504866623 | *Methanolobus psychrophilus* | 641 | MBL fold hydrolase | 30.77 | 1.73E-05 | 51.17 |
| gi\|816389003 | *Lokiarchaeum* sp. GC14_75 | 648 | Metallo-beta-lactamase L1 | 30.77 | 3.64E-08 | 59.62 |
| gi\|851262085 | *Methanosarcina horonobensis* | 641 | MBL fold hydrolase | 30.77 | 1.56E-04 | 51.17 |
| gi\|502745672 | *Methanocaldococcus* sp. FS406-22 | 558 | MBL fold metallo-hydrolase | 30.70 | 5.98E-05 | 50.70 |
| gi\|170934313 | *Pyrobaculum neutrophilum* V24Sta | 615 | Beta-lactamase domain protein | 30.65 | 1.60E-06 | 53.05 |
| gi\|757124828 | *Thermococcus paralvinellae* | 717 | Zn-dependent hydrolase | 30.61 | 0.002 | 55.87 |
| gi\|756792592 | *Candidatus Nitrosopumilus piranensis* | 1404 | Rhodanese domain-containing protein | 30.56 | 2.45E-05 | 63.38 |
| gi\|851287001 | *Palaeococcus ferrophilus* | 609 | Glyoxalase | 30.51 | 8.05E-04 | 50.70 |
| gi\|973113610 | *Methanocalculus* sp. 52_23 | 603 | Beta-lactamase domain protein | 30.40 | 8.24E-06 | 51.17 |
| gi\|973162189 | *Methanomicrobiales archaeon* 53_19 | 603 | Beta-lactamase domain protein | 30.40 | 6.11E-06 | 51.17 |
| gi\|524837456 | *Methanoculleus* sp. CAG:1088 | 641 | Putative uncharacterized protein | 30.37 | 0.035 | 50.70 |
| gi\|851257745 | *Methanosarcina* | 600 | Hypothetical protein | 30.37 | 0.13 | 59.62 |
| gi\|700303882 | *Thermococcus eurythermalis* | 681 | Hydrolase | 30.28 | 2.08E-04 | 52.11 |
| gi\|735015437 | archaeon GW2011_AR11 | 714 | Beta-lactamase protein | 30.25 | 0.23 | 50.70 |
| gi\|15623131 | *Sulfolobus tokodaii* str. 7 | 600 | Putative hydrolase | 30.23 | 7.15E-06 | 54.46 |
| gi\|500271928 | *Metallosphaera sedula* | 606 | MBL fold metallo-hydrolase | 30.23 | 6.05E-04 | 53.52 |
| gi\|919520712 | *Sulfolobus tokodaii* | 597 | Hypothetical protein | 30.23 | 6.99E-06 | 54.46 |

| | | | | | | | |
|---|---|---|---|---|---|---|---|
| | gi\|851163782 | *Geoglobus acetivorans* | 615 | Zn-dependent hydrolase | 30.17 | 1.29E-04 | 50.70 |
| | gi\|851219309 | *Candidatus Methanoplasma termitum* | 576 | MBL fold metallo-hydrolase | 30.08 | 0.032 | 52.58 |
| | gi\|494104154 | *Methanotorris formicicus* | 618 | Hypothetical protein | 30.08 | 4.44E-06 | 53.05 |
| | gi\|329138039 | *Candidatus Nitrosoarchaeum limnia* | 1398 | Rhodanese domain-containing protein | 30.07 | 6.43E-07 | 63.38 |
| | gi\|500766631 | *Methanoregula boonei* | 696 | MBL fold metallo-hydrolase | 30.07 | 0.055 | 56.34 |
| **Ancestor class C β-lactamase** | gi\|445600523 | *Natronococcus amylolyticus* DSM10524 | 1974 | Beta-lactamase | 31.07 | 1.66E-13 | 52.02 |
| | gi\|1011504614 | *Methanogenium cariaci* | 834 | Hypothetical protein | 30.80 | 7.49E-12 | 57.68 |
| | **gi\|919167542** | ***Methanosarcina sp.* WH1** | **1959** | **Hypothetical protein** | **30.63** | **1.45E-17** | **75.74** |
| | gi\|919167113 | *Methanosarcina sp.* WWM596 | 1959 | Hypothetical protein | 30.63 | 1.82E-17 | 75.74 |

**Suppl. Table S2:** Three-dimensional (3D) structure comparison with available and characterized proteins from the Phyre$^2$ investigator database.

| **Archaea** | | **Phyre$^2$ investigator database** | | | | | |
|---|---|---|---|---|---|---|---|
| **Protein sequences** | **Size (aa)** | **Best protein hit** | **Hit ID** | **Confidence** | **Coverage** | **Comments** | **3D Structure** |
| **Class B β-lactamase** | **213** | Crystal structure of New Delhi Metallo-β-lactamase (NDM-1) form *Klebsiella pneumoniae* | c3rkjA | 100% | 94% | 201 residues (94%) have been modelled with 100% confidence by the single highest scoring template | 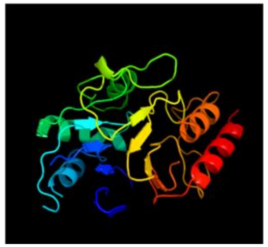 |
| **Class C-like β-lactamase** | **653** | Structure of the octameric penicillin-binding protein (PBP) homologue from *pyrococcus abyssi* | c2qmiH | 100% | 66% | 430 residues (64%) have been modelled with 100% confidence by the single highest scoring template | 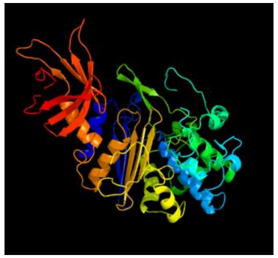 |

**Suppl. Table S3**: Antibiotic resistance pattern of *Methanosarcina* compared to *Elizabethkingia* GOB-15

| | ***M. barkeri* class B** | ***Methanosarcina* sp. class C-like** | ***Elizabethkingia* GOB-13** [13] |
|---|---|---|---|
| *β*-lactams | | | |
| Ampicillin | R | R | R |
| Ampicillin/sulbactam | R | R | R |
| Penicillin | R | R | R |
| Piperacillin | R | R | R |
| Piperacillin/tazobactam | R | R | R |
| Cefoxitin | R | R | R |
| Ceftriaxone | R | R | R |
| Ceftazidime | R | R | R |
| Imipenem | R | R | R |
| Meropenem | R | R | R |
| Aztreonam | R | R | R |
| Non *β*-lactams | | | |
| Gentamicin | R | R | S |
| Ciprofloxacin | R | R | R |
| Amikacin | R | R | S |
| Trimethoprim-sulfamethoxazole | R | R | R |

Nd - not determined. R, resistance; S, Susceptible.

13 - Opota et al., IJAA 49, 93-97 (2016)

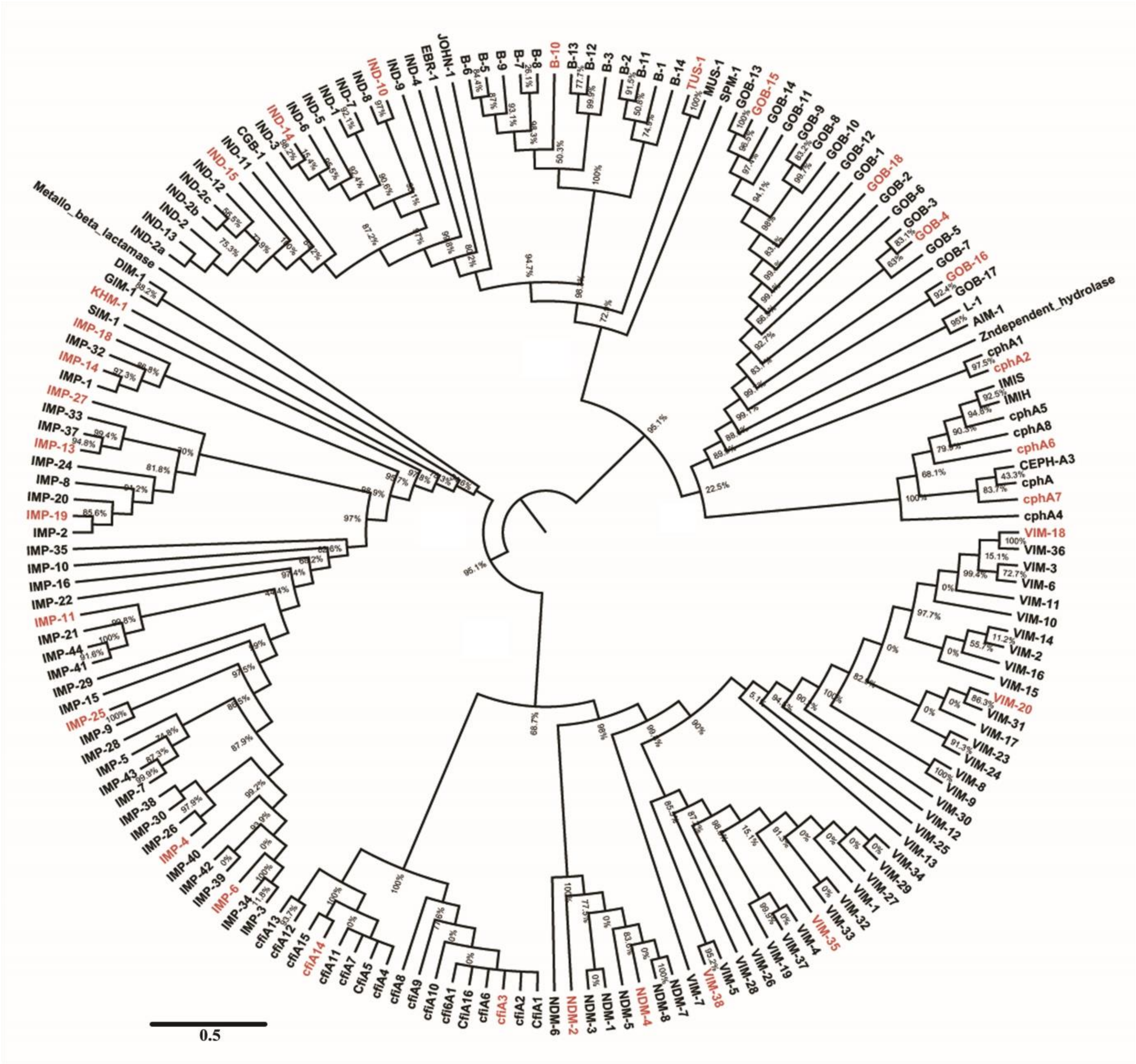

Metallo_beta_lactamase
IND-2a
IND-13
IND-2
IND-2b
IND-2c
IND-12
IND-15
IND-11
CGB-1
IND-3
IND-14
IND-6
IND-5
IND-1
IND-7
IND-8
IND-10
IND-9
IND-4
EBR-1
JOHN-1
B-6
B-5
B-9
B-7
B-8
B-10
B-13
B-12
B-3
B-2
B-11
B-1
B-14
TUS-1
MUS-1
SPM-1
GOB-13
GOB-15
GOB-14
GOB-11
GOB-9
GOB-8
GOB-10
GOB-12
GOB-1
GOB-18
GOB-2
GOB-6
GOB-3
GOB-4
GOB-5
GOB-7
GOB-16
GOB-17
L-1
AIM-1
Zndependent_hydrolase
cphA1
cphA2
IMIS
IMIH
cphA5
cphA8
cphA6
CEPH-A3
cphA
cphA7
cphA4
VIM-18
VIM-36
VIM-3
VIM-6
VIM-11
VIM-10
VIM-14
VIM-2
VIM-16
VIM-15
VIM-20
VIM-31
VIM-17
VIM-23
VIM-24
VIM-8
VIM-9
VIM-30
VIM-12
VIM-25
VIM-13
VIM-34
VIM-29
VIM-27
VIM-1
VIM-32
VIM-33
VIM-35
VIM-4
VIM-37
VIM-19
VIM-26
VIM-28
VIM-5
VIM-38
VIM-7
NDM-7
NDM-8
NDM-4
NDM-5
NDM-1
NDM-3
NDM-2
NDM-6
CfiA1
cfiA2
cfiA3
cfiA6
CfiA16
cfi6A1
cfiA10
cfiA9
cfiA8
cfiA4
CfiA5
cfiA7
cfiA11
cfiA14
cfiA15
cfiA12
cfiA13
IMP-3
IMP-34
IMP-6
IMP-39
IMP-42
IMP-40
IMP-4
IMP-26
IMP-30
IMP-38
IMP-7
IMP-43
IMP-5
IMP-28
IMP-9
IMP-25
IMP-15
IMP-29
IMP-41
IMP-44
IMP-21
IMP-11
IMP-22
IMP-16
IMP-10
IMP-35
IMP-2
IMP-19
IMP-20
IMP-8
IMP-24
IMP-13
IMP-37
IMP-33
IMP-27
IMP-1
IMP-14
IMP-32
IMP-18
SIM-1
KHM-1
GIM-1
DIM-1
0.5

**Suppl. Figure S1: Phylogenetic Tree of Class B β-lactamases: The phylogenetic tree was inferred using the** approximate Maximum Likelihood method under the JTT matrix-based model. The analysis involved 174 amino acid sequences from class B β-lactamases. Evolutionary analysis was conducted in FastTree and visualized in FigTree. The existing clades are labeled as b-1 to b-4.

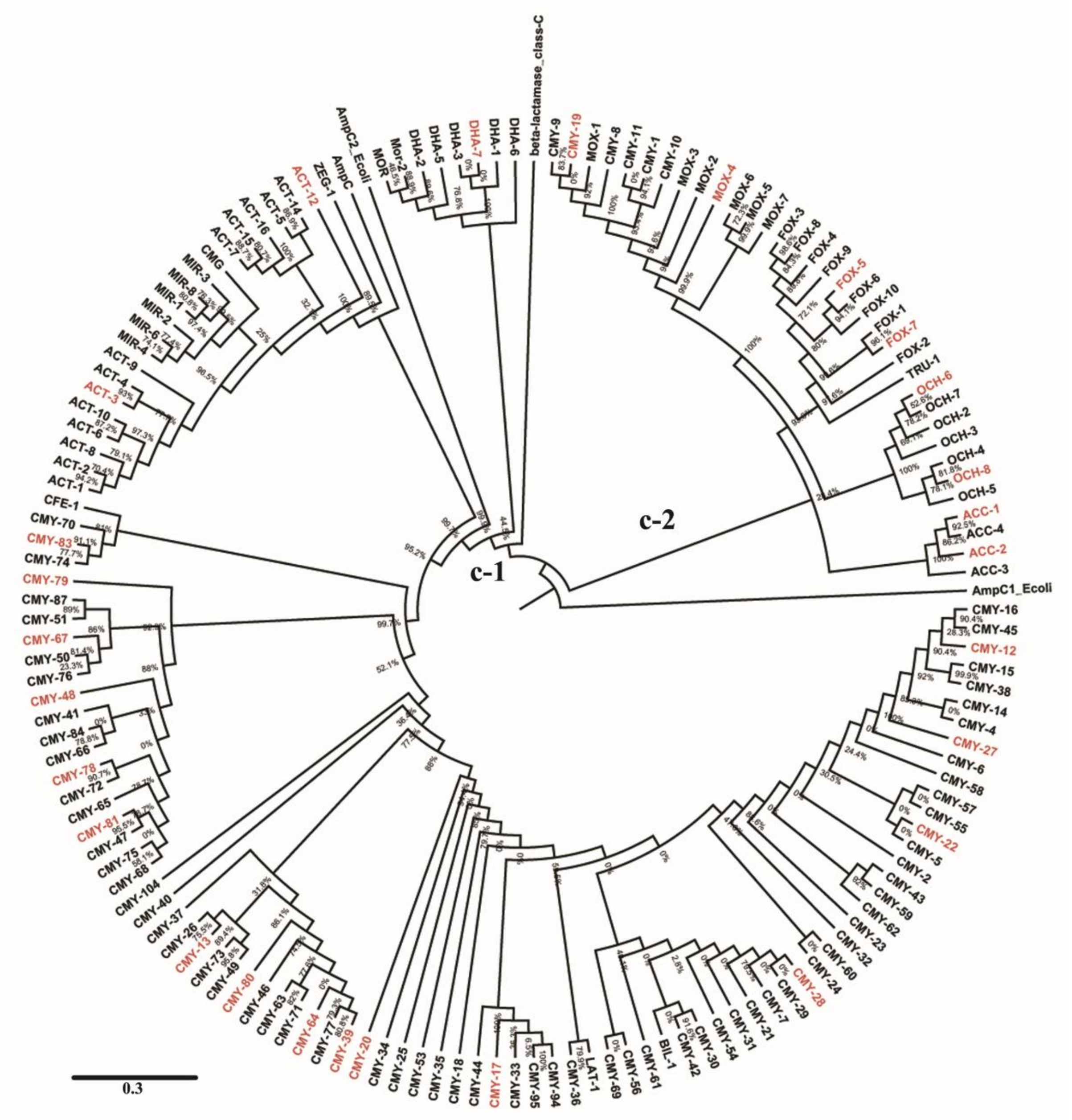
beta-lactamase_class-C
DHA-6
DHA-1
DHA-7
DHA-3
DHA-5
DHA-2
Mor-2
MOR
AmpC2_Ecoli
AmpC
ZEG-1
ACT-12
ACT-14
ACT-5
ACT-16
ACT-15
ACT-7
CMG
MIR-3
MIR-8
MIR-1
MIR-2
MIR-6
MIR-4
ACT-9
ACT-4
ACT-3
ACT-10
ACT-6
ACT-8
ACT-2
ACT-1
CFE-1
CMY-70
CMY-83
CMY-74
CMY-79
CMY-87
CMY-51
CMY-67
CMY-50
CMY-76
CMY-48
CMY-41
CMY-84
CMY-66
CMY-78
CMY-72
CMY-65
CMY-81
CMY-47
CMY-75
CMY-68
CMY-104
CMY-40
CMY-37
CMY-26
CMY-13
CMY-73
CMY-49
CMY-80
CMY-46
CMY-63
CMY-71
CMY-64
CMY-77
CMY-39
CMY-20
CMY-34
CMY-25
CMY-53
CMY-35
CMY-18
CMY-44
CMY-17
CMY-33
CMY-95
CMY-94
CMY-36
LAT-1
CMY-69
CMY-56
CMY-61
BIL-1
CMY-42
CMY-30
CMY-54
CMY-31
CMY-21
CMY-7
CMY-29
CMY-28
CMY-24
CMY-60
CMY-32
CMY-23
CMY-62
CMY-59
CMY-43
CMY-2
CMY-5
CMY-22
CMY-55
CMY-57
CMY-58
CMY-6
CMY-27
CMY-4
CMY-14
CMY-38
CMY-15
CMY-12
CMY-45
CMY-16
AmpC1_Ecoli
ACC-3
ACC-2
ACC-4
ACC-1
OCH-5
OCH-8
OCH-4
OCH-3
OCH-2
OCH-7
OCH-6
TRU-1
FOX-2
FOX-7
FOX-1
FOX-10
FOX-6
FOX-5
FOX-9
FOX-4
FOX-8
FOX-3
MOX-7
MOX-5
MOX-6
MOX-4
MOX-2
MOX-3
CMY-10
CMY-1
CMY-11
CMY-8
MOX-1
CMY-19
CMY-9
c-1
c-2
0.3

**Suppl. Figure S2: Phylogenetic tree of Class C-like β-lactamases: The phylogenetic tree was inferred using the** approximate Maximum Likelihood method under the JTT matrix-based model. The analysis involved 151 amino acid sequences from the class C-like β-lactamases. Evolutionary analysis was conducted in FastTree and visualized in FigTree. The existing clades are labeled as c-1 and c-2.

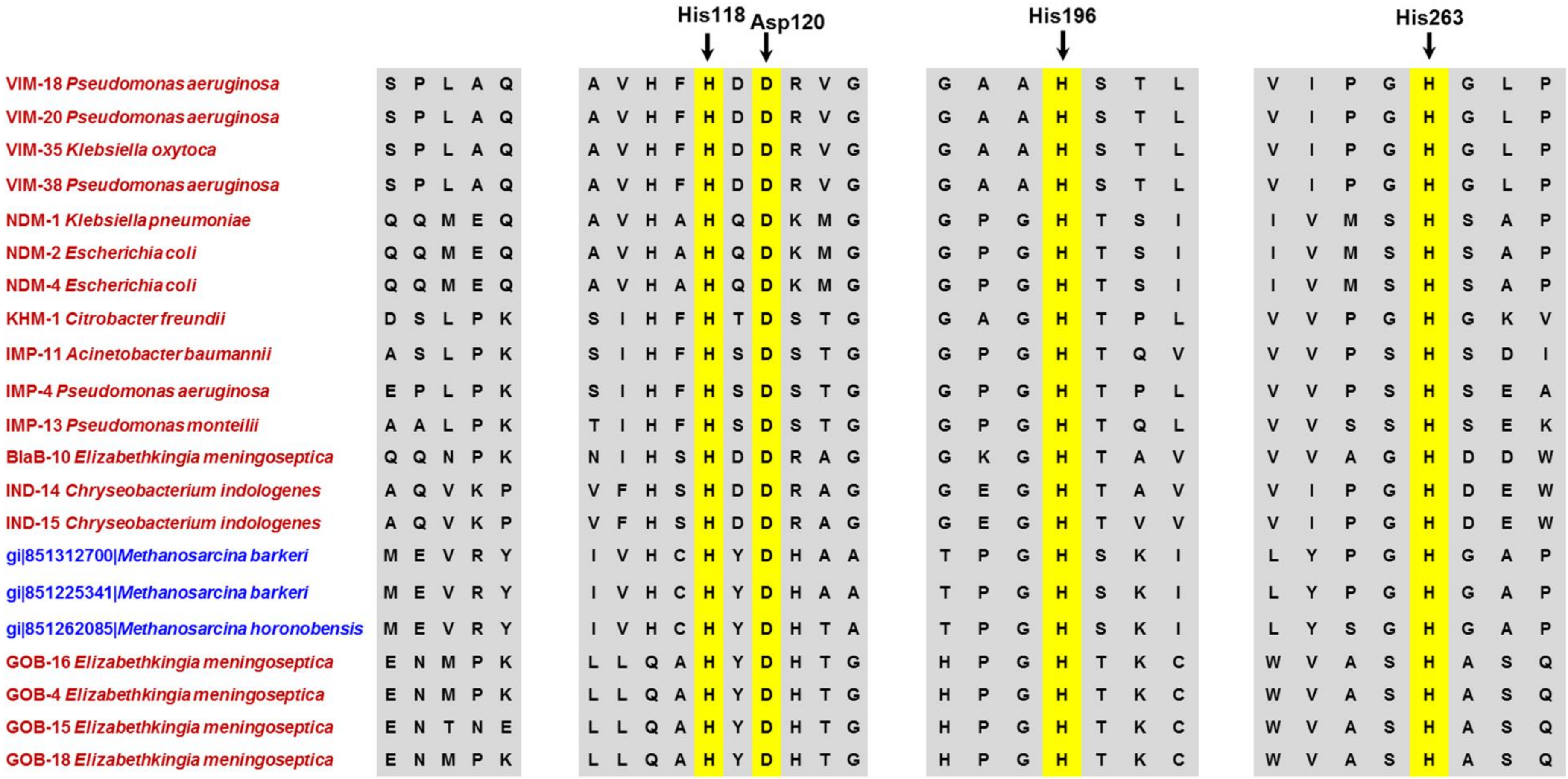


**Suppl. Figure S3**: Protein sequences alignment of bacterial and archaeal class B β-lactamases. Known and conserved residues of bacterial metallo-β-lactamases are highlighted with yellow color.

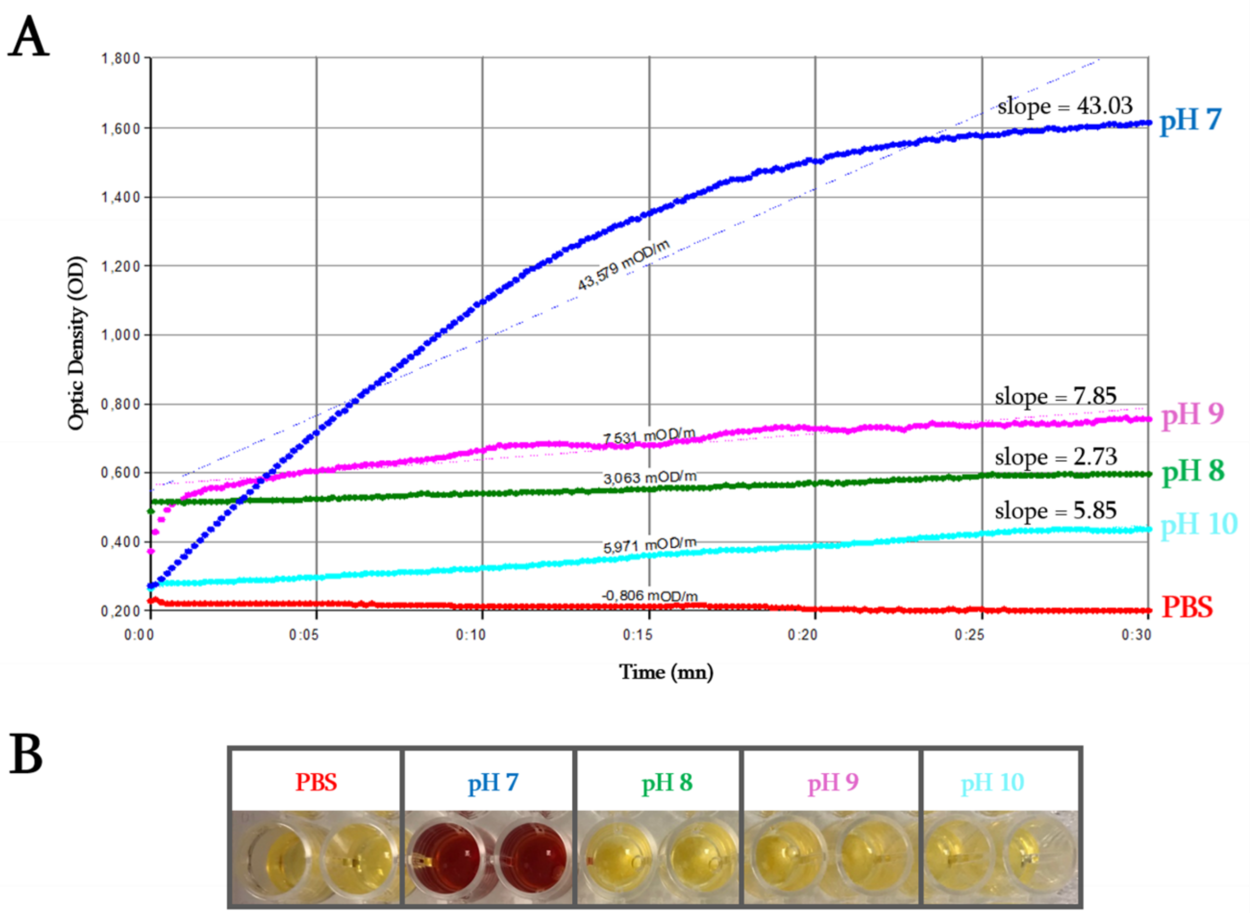


**Suppl. Figure S4:** Evaluation of the archaeal MetbaB enzyme activity on nitrocefin at different pH. **(A)** The monitoring of the nitrocefin degradation by MetbaB enzymes during 30 minutes under the four different pH conditions. **(B)** The MetbaB activity test on the chromogenic cephalosporin substrate in liquid medium.

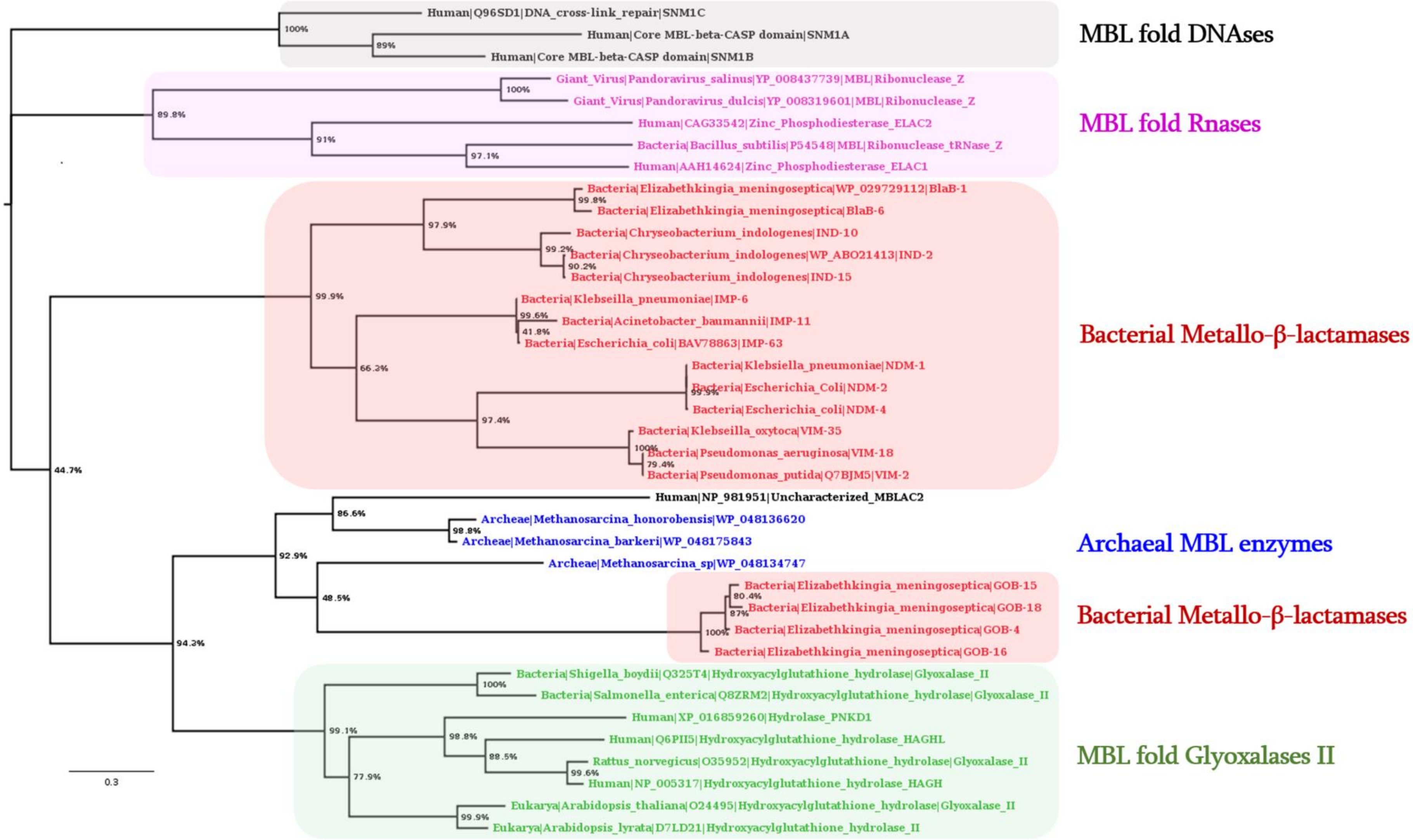

Human|Q96SD1|DNA_cross-link_repair|SNM1C
Human|Core MBL-beta-CASP domain|SNM1A
Human|Core MBL-beta-CASP domain|SNM1B
MBL fold DNAses
Giant_Virus|Pandoravirus_salinus|YP_008437739|MBL|Ribonuclease_Z
Giant_Virus|Pandoravirus_dulcis|YP_008319601|MBL|Ribonuclease_Z
Human|CAG33542|Zinc_Phosphodiesterase_ELAC2
Bacteria|Bacillus_subtilis|P54548|MBL|Ribonuclease_tRNase_Z
Human|AAH14624|Zinc_Phosphodiesterase_ELAC1
MBL fold Rnases
Bacteria|Elizabethkingia_meningoseptica|WP_029729112|BlaB-1
Bacteria|Elizabethkingia_meningoseptica|BlaB-6
Bacteria|Chryseobacterium_indologenes|IND-10
Bacteria|Chryseobacterium_indologenes|WP_ABO21413|IND-2
Bacteria|Chryseobacterium_indologenes|IND-15
Bacteria|Klebseilla_pneumoniae|IMP-6
Bacteria|Acinetobacter_baumannii|IMP-11
Bacteria|Escherichia_coli|BAV78863|IMP-63
Bacteria|Klebsiella_pneumoniae|NDM-1
Bacteria|Escherichia_Coli|NDM-2
Bacteria|Escherichia_coli|NDM-4
Bacteria|Klebseilla_oxytoca|VIM-35
Bacteria|Pseudomonas_aeruginosa|VIM-18
Bacteria|Pseudomonas_putida|Q7BJM5|VIM-2
Bacterial Metallo-β-lactamases
Human|NP_981951|Uncharacterized_MBLAC2
Archeae|Methanosarcina_honorobensis|WP_048136620
Archeae|Methanosarcina_barkeri|WP_048175843
Archeae|Methanosarcina_sp|WP_048134747
Archaeal MBL enzymes
Bacteria|Elizabethkingia_meningoseptica|GOB-15
Bacteria|Elizabethkingia_meningoseptica|GOB-18
Bacteria|Elizabethkingia_meningoseptica|GOB-4
Bacteria|Elizabethkingia_meningoseptica|GOB-16
Bacterial Metallo-β-lactamases
Bacteria|Shigella_boydii|Q325T4|Hydroxyacylglutathione_hydrolase|Glyoxalase_II
Bacteria|Salmonella_enterica|Q8ZRM2|Hydroxyacylglutathione_hydrolase|Glyoxalase_II
Human|XP_016859260|Hydrolase_PNKD1
Human|Q6PII5|Hydroxyacylglutathione_hydrolase_HAGHL
Rattus_norvegicus|O35952|Hydroxyacylglutathione_hydrolase|Glyoxalase_II
Human|NP_005317|Hydroxyacylglutathione_hydrolase_HAGH
Eukarya|Arabidopsis_thaliana|O24495|Hydroxyacylglutathione_hydrolase|Glyoxalase_II
Eukarya|Arabidopsis_lyrata|D7LD21|Hydroxyacylglutathione_hydrolase_II
MBL fold Glyoxalases II
0.3

**Suppl. Figure S5:** Phylogenetic relation cheap of different MBL fold proteins including β-lactamases, nucleases, ribonucleases, glyoxalase, and identified enzymes from human, bacteria, archaea, and giant virus.

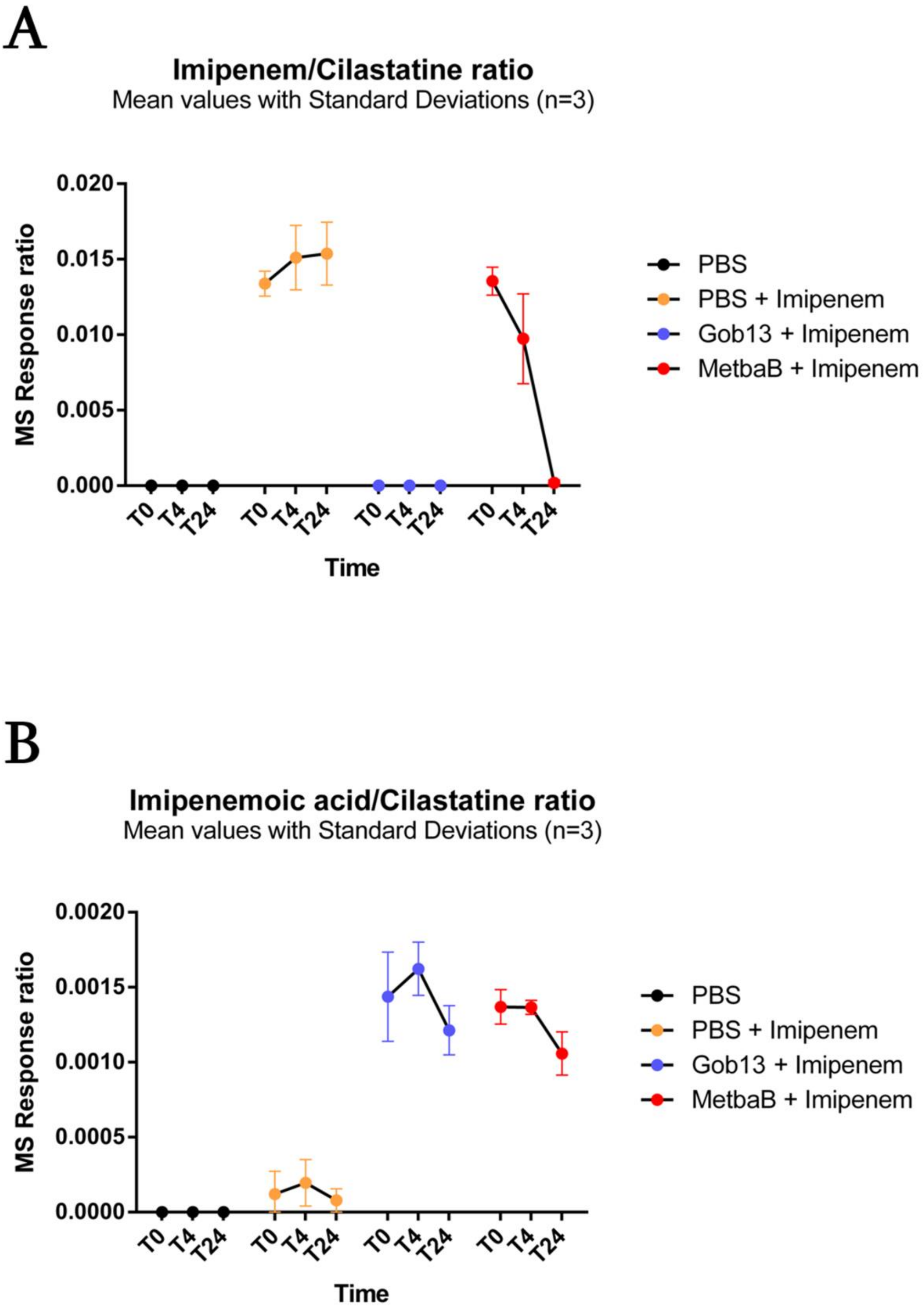


**Suppl. Figure S6:** Monitoring the imipenem hydrolysis by the *E. meningoseptica* metallo-β-lactamase GOB-13 and MetbaB enzyme by LC-MS. Time depended production of the metabolite of imipenem (i.e. imipenemoic acid) is measured here using a unhydrolyzed cilastatine substrate. Both enzymes hydrolyze efficiently imipenem through the increase accumulation of its metabolite.

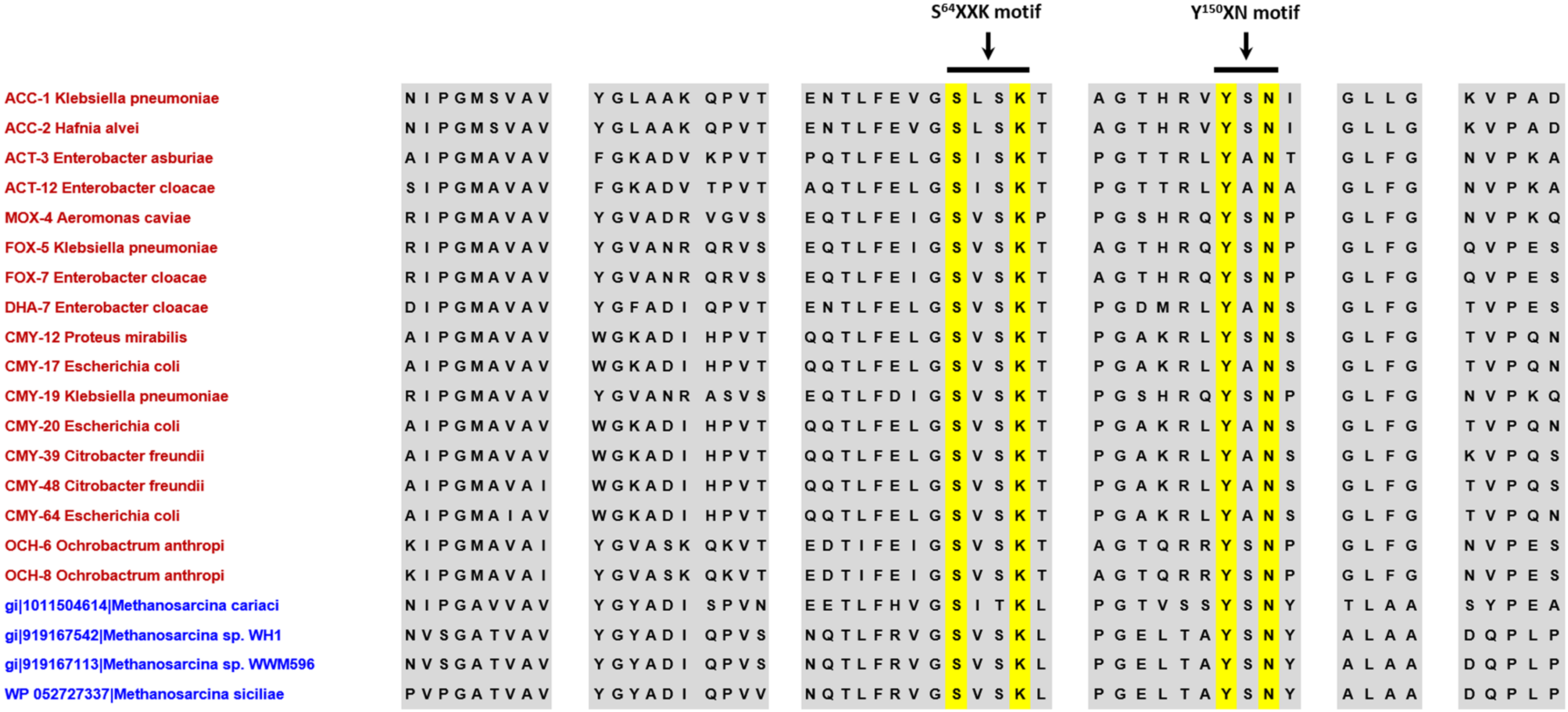


**Suppl. Figure S7**: Protein alignment sequences of bacterial and archaeal class C-like β-lactamase proteins. Known and conserved motifs of bacterial class C β-lactamases are highlighted with yellow color.

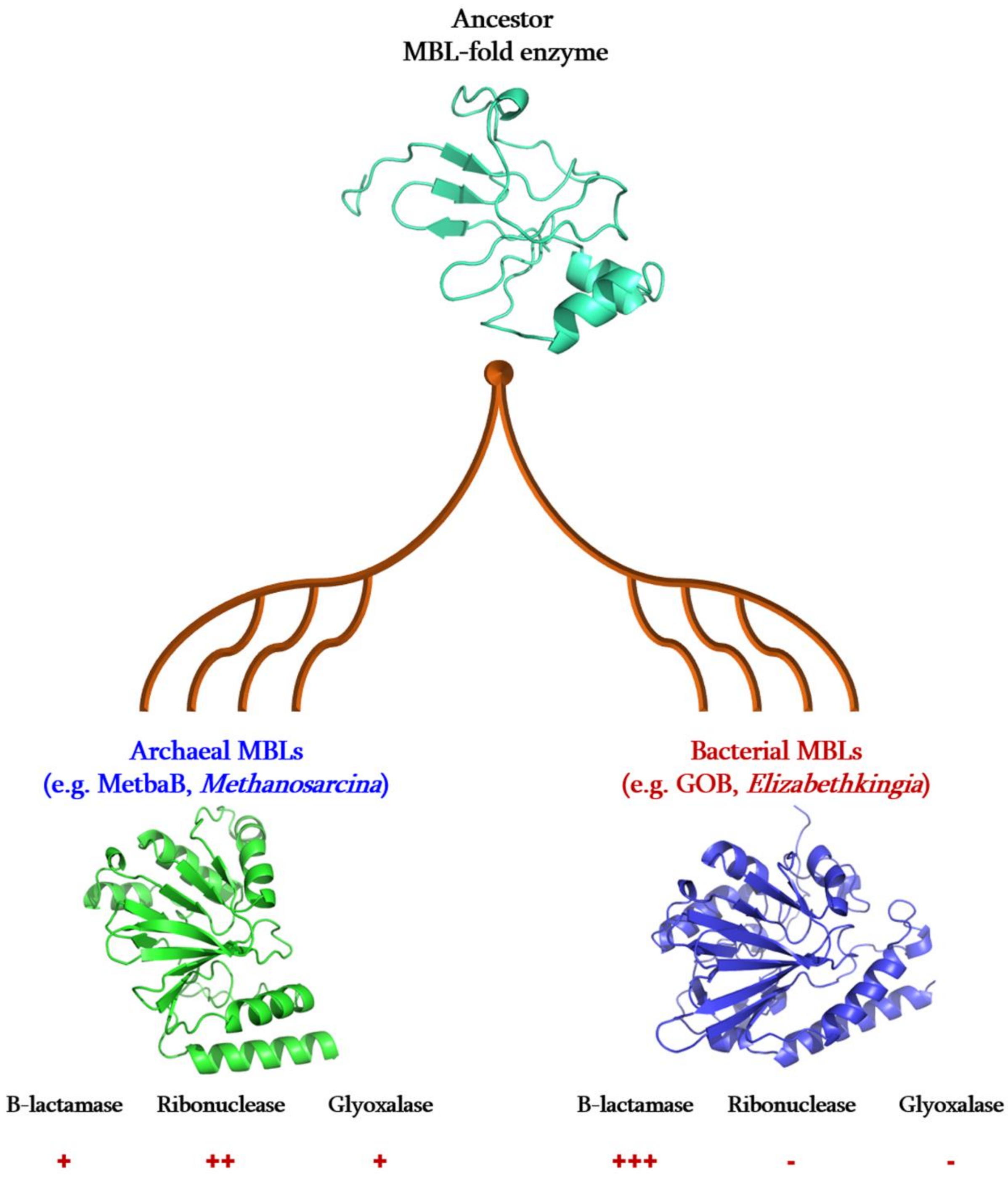


**Suppl. Figure S8:** Putative evolution scenario of enzymatic activities of MBL fold proteins.

(+) : positive activity; (-) : negative activity.
